# Early life adversity disrupts adult social reward motivation

**DOI:** 10.64898/2026.07.21.739864

**Authors:** Margie C. Stricklin, Judy E. Nyakoa, Kevin M. Mesape, Moushmi S. Bhat, Siarra S. Rolle, Debra A. Bangasser, Amelia Cuarenta

**Affiliations:** Neuroscience Institute, Georgia State University, Atlanta, GA, USA; Center for Behavioral Neuroscience, Atlanta, GA, USA; Department of Psychology, University of Michigan, Ann Arbor, MI, USA

## Abstract

Early life adversity can produce persistent changes during development that increase vulnerability to neuropsychiatric disorders. Although disruptions in reward processing are widely recognized as a hallmark of these disorders, reward is not a single construct. Distinct forms of reward including social interaction and primary rewards such as food rely on overlapping but dissociable neural circuitry and may be differentially affected by adverse experiences. Here, we used a rodent model of neonatal predator odor exposure (POE) to determine how early life threat influences motivation for social and sucrose reward in adulthood using operant procedures. Neonatal POE reduced adult motivation for a social reinforcer during an operant social self-administration task, whereas motivation for a sucrose reinforcer was unchanged. However, we did find a significant difference in sucrose self-administration with POE females pressing more for sucrose than control females. These findings demonstrate that neonatal threat does not produce a generalized deficit in motivation but rather selectively alters motivation across distinct reward domains. Together, this work identifies social reward as a particularly vulnerable behavioral domain following early life threat and provides new insight into how adverse developmental experiences shape adult reward-related behavior.

## Introduction

Early life adversity (ELA) is a common environmental experience strongly associated with an increased risk for a range of psychiatric disorders across the lifespan, including depression, anxiety, and substance use disorders^1-4^. A substantial proportion of individuals experience at least one form of adversity during childhood, such as abuse, neglect, or household dysfunction^5-8^, underscoring its broad public health relevance. Early life represents a sensitive developmental window during which environmental experiences can have profound and lasting effects on the brain and behavior. Adverse experiences during this period, including exposure to threat or instability, have been shown to shape the maturation of neural systems involved in emotion, motivation, and social behavior, with impacts extending into adulthood^9-11^. Converging evidence from both human and animal studies suggests that ELA is associated with alterations in reward processing and social functioning^12-16^. However, the extent to which specific features of early experiences, such as exposure to threat, produce enduring versus transient changes in these systems remains unclear.

Social behavior is a critical component of development, supporting the formation of social bonds, the acquisition of adaptive behavioral strategies, and the regulation of emotional processes across the lifespan^17,18^. In rodents, social interactions such as juvenile play behavior represent a highly conserved and developmentally sensitive form of social engagement that emerges early and is essential for the maturation of social and affect-related processes and cognitive functions^17,19,20^. Disruptions in social behavior during development have been associated with persistent changes in affective and anxiety-like behavioral outcomes^21-23^. Importantly, social behavior is particularly sensitive to environmental input during early life, as it depends on the coordinated maturation of limbic and stress-responsive circuits that are actively developing during this period.

Our laboratory utilizes a predator odor exposure (POE) model in rats to mimic early-life threat in which pups are briefly exposed to predator odors during the early neonatal period. Predator odors represent ethologically relevant threat cues that engage stress-responsive systems^24-26^ without direct physical harm, providing a naturalistic approach for examining the effects of early-life threat exposure. This developmental period coincides with rapid maturation of limbic and stress-responsive systems^27,28^, making it particularly sensitive to environmental input. Using this model, we have previously found a reduction in social play and an increase in anxiety-like behavior during the juvenile stage of development^29,30^. However, it remains unclear whether early-life POE produces enduring behavioral changes into adulthood or instead reflect transient developmental adaptations.

Here, we address this gap by determining whether neonatal POE produces lasting alterations in social behaviors and reward motivation more broadly. Social motivation was assessed using a social self-administration paradigm, an operant task that uses social interaction as a reinforcer. In this task, rats are trained to perform an operant response that results in the opening of a door and a brief opportunity to interact with a same-sex conspecific housed in an adjacent chamber. By requiring animals to actively work for social contact, this approach allows for assessment of the reinforcing and motivational properties of social reward, and the effort animals are willing to expend to obtain a social interaction. Sucrose self-administration was included to assess motivation for another form of natural reward and to determine whether the effects of neonatal POE generalized across reward domains or was selective for a specific type of reward.

Based on our prior evidence of reduced juvenile play behavior following neonatal POE, we hypothesized that early life threat exposure would reorganize motivational priorities in adulthood. Specifically, we predicted that POE would be associated with altered sucrose self-administration and reduced social motivation, reflecting a shift in reward valuation following ELA. These studies are designed to fill a critical gap by linking early life behavioral changes to adult outcomes thereby providing insight into the long-term consequences of early life threat exposure on reward processing.

## Methods

### Subjects and predator odor exposure (POE)

Georgia State University Institutional Animal Care and Use Committee approved all animal care and experiments in this study. Long Evans rats were obtained from Charles River Laboratories (Wilmington, MA) and bred in-house with their offspring used for studies. All POE and control dams were primiparous. For all experiments, no more than two males and two females (sex defined by external genitalia) were used from each litter, and pups were randomly assigned to experiments. On postnatal day (PND) 1, litters were culled to ten pups with 6M/4F or 6F/4M composition; one litter contained only nine pups (5M/4F) because the dam had only nine offspring. Neonatal rat pups were briefly removed from their dam (<15 min) and brought to a separate room where they were placed on an elevated platform in an enclosure containing bedding (Bed-o’Cobs) mixed with predator odor; pups did not have direct contact with the odor-infused bedding. The exposure lasted for 5 minutes and occurred on 3 consecutive days. On PND1 rat pups were exposed to a bobcat odor, on PND2 rat pups were exposed to an unrelated novel adult male rat odor, and on PND3 rat pups were exposed to a ferret odor. Different odors were used each day to avoid odor habituation. Pups were monitored throughout the entire exposure period. Animals in the control group were handled in the same manner but instead of being exposed to bedding mixed with predator odor they were exposed to clean bedding (Bed-o’Cobs, The Andersons) only. All animals were returned to the dam in <15 minutes. Rats were weaned on PND 21 and pair-housed with rats of the same sex and condition (i.e., POE vs. control housing). On PND 60, animals were handled daily prior to experimental testing beginning on PND 70. Throughout the study, rats were housed in a reverse dark/light cycle with lights on from 9:00PM to 9:00AM with behavioral studies conducted between 11:00AM-2:00PM. All rats had *ad libitum* access to water and standard laboratory chow (5001, LabDiet) throughout the duration of the experiment. All housing occurred within standard shoebox cages (20×20×40 cm) with vent filter cage tops (on non-ventilated racks). No rats were dropped from these experiments. All rats were run on sucrose self-administration first, followed by social self-administration.

### Sucrose self-administration

Male and female rats performed oral self-administration using MedPC Operant Chambers. Behavioral studies occurred during the dark phase. Behavioral consisted of a 1-hour self-administration session in the dark cycle. Rats were tested under satiated conditions to minimize the influence of metabolic need on motivated responding for sucrose reward. Control and POE rats were first trained for 4 days to respond for sucrose reward on a fixed ratio (FR) 1 schedule, where one active lever press resulted in one sucrose pellet reward paired with a 5-s cue light. During this phase, rats were able to receive a maximum of 90 sucrose pellets per session. Following 4 days of FR1, response requirements were increased in a stepwise manner across FR schedules to strengthen operant responding prior to PR testing. Specifically, rats completed 2 days each under FR2, FR4, and FR8 schedules, in which 2, 4, or 8 active lever presses, respectively, were required to receive a single sucrose pellet. Rats were then tested for motivation under a progressive ratio (PR) schedule during a single session, where the number of lever presses required to earn each subsequent social interaction increased over each trial. Under a PR schedule, the response requirement for each subsequent sucrose pellet delivery increased exponentially until the subject failed to meet the requirement. The response requirement of the *i*th reinforcement was given by R(*i*)=[5e^0.2i^-5], and the session expired when an animal took more than 60 min to earn a sucrose pellet^31-33^.

### Social self-administration

Rats were trained to self-administer for access to a same-sex age-matched conspecific (“target” rat) raised under control housing conditions during daily 60-min sessions in social motivation chambers (Med Associates; ENV-008CT). Each target rat was used for two consecutive days before being replaced with a novel target. Trials began with the illumination of a white house light, and 10 s later, a social-paired active lever released. Rats were given 60 s to press the active lever on an FR1 reinforcement schedule before it automatically retracted, and the house light turned off. Pressing the active lever resulted in retraction of the lever and the lifting of a guillotine-style door. After the door lifted, the resident rat was able to interact with the target rat for 40 s. A barrier was present keeping the rats from going into each other’s chamber, but they were able to interact via sniffing and touching. After this period, the house light turned off, the guillotine door closed, and a new trial began shortly thereafter. After 4 days of FR1, rats were trained on 2 days of FR2, FR4, and FR8 each before being tested on a progressive ratio (PR) schedule. During the progressive ratio sessions, we increased the ratio of responses per reward as detailed^34^. The final completed response ratio represents the “breakpoint” value.

### Statistical analysis

Sample sizes were informed by prior studies using similar behavioral procedures and outcomes^31,35-37^. Final sample sizes are reported in each figure legend and reflect the number of animals that completed testing and met prespecified inclusion criteria. Exclusions due to failure to meet acquisition criteria is reported where applicable. All statistical analyses were performed in GraphPad Prism. All significance thresholds were set at *p*<.05.

## Results

### Effects of early life threat on sucrose motivation

To determine whether early life threat alters motivation for sucrose, male and female control and POE rats were assessed in an operant sucrose self-administration task. Sucrose-reinforced responding was assessed under fixed-ratio schedules, followed by a PR test of motivation. There was no significant 3-way Condition × Sex × Day interaction (*F*_3,135_ < 1.0); however, we did find a significant Condition × Sex interaction in active lever presses (*F*_1,45_ = 4.098, *p*<.05), indicating that the effect of POE on sucrose self-administration differed between males and females (Fig. 1A & 1B). No significant day × sex (F_3,135_ <1.0, *p*=.72) or day × condition (F_3,135_ <1.0, *p*=.59) interactions were observed in active lever presses. There were no main effects of sex (F_1,45_ = 1.900, *p*=.17) or condition (F_1,45_ = 1.962, *p*=.17). There was a significant impact of day on active lever presses, (F_3,135_ = 32.61, *p*<.001), indicating that the number of active lever presses differed across days.

**Figure 1.**
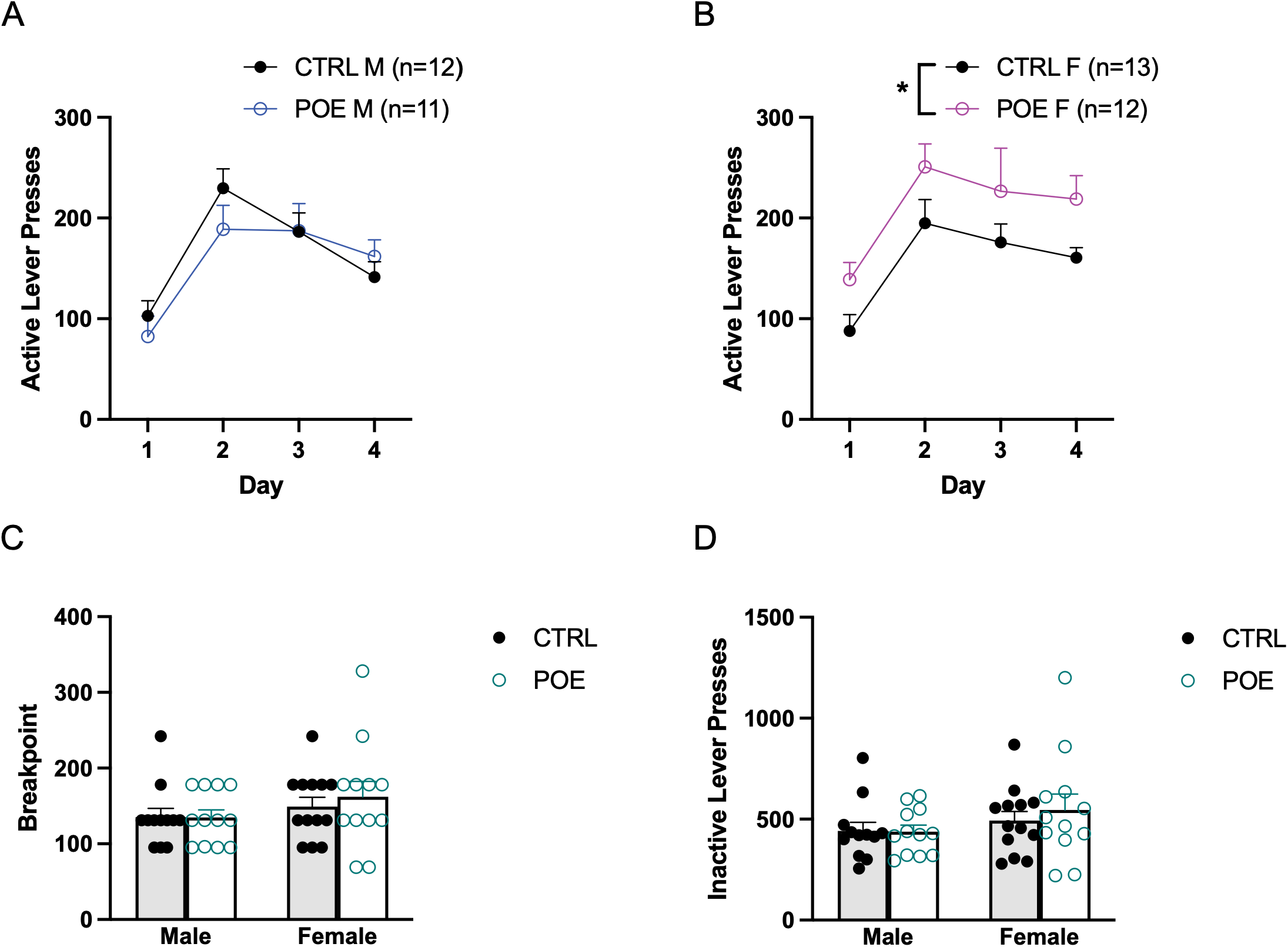
Effects of POE on sucrose self-administration. A) Number of active lever presses on an FR1 schedule over 4 days in males. B) Number of active lever presses on an FR1 schedule over 4 days in females. C) Breakpoint ratio on a PR schedule. D) Number of active lever presses on a PR schedule. N=11-13/group. * *p* <.05. Data are represented as means ± SEMs. CTRL, control; POE, Predator Odor Exposure; M, Male; F, Female.

There was no significant 3-way Condition × Sex × Day interaction (*F*_2.58,116.4_ = 1.373, *p*=.26) for rewards. No significant day × sex (F_2.87,116.4_ = .6959, *p*=.54) or day × condition (F_2.587,116.4_ <1.0, *p*=.76) interactions were observed in rewards obtained. There were no main effects of sex (F_1,45_ <1.0, *p*=.63) or condition (F_1,45_ <1.0, *p*=.69). There was a significant impact of day on rewards (F_2.587,116.4_ = 100.2, *p*<.001), indicating that the number of rewards differed across days.

Rats were tested under a PR schedule to assess sucrose motivation. A two-way ANOVA revealed no significant Condition × Sex interactions on rewards earned (F_1,45_ =<1.0, *p*=.86), active lever presses (F_1,45_ <1.0, *p*=.59), breakpoint (F_1,45_ <1.0, *p*=.65), or inactive lever presses (F_1,45_ =1.723, *p*=.20), nor were there any significant effects of condition on rewards earned (*F*_1,45_<1.0, *p*=.86), active lever presses (*F*_1,45_ <1.0, *p*=.65), breakpoint (*F*_1,45_ <1.0, *p*=.67), or inactive lever presses (*F*_1,45_ <1.0, *p*=.39) (Fig 1C and 1D). There was also no significant sex effects on rewards earned (F_1,45_ =1.454, *p*=.23), active lever presses (F_1,45_ =2.276, *p*=.14), breakpoint (F_1,45_ =2.124, *p*=.15), or inactive lever presses (F_1,45_ =1.016, *p*=.32).

### Effects of early life threat on social motivation

To test the effects of early life threat on motivation for social rewards, we compared POE males and females to control males and females in an operant task in which animals responded for access to a novel same-sex and age-matched conspecific. We conducted a 3-way ANOVA and found no significant Condition × Sex × Day interaction (*F*_2.386,105_ = 0.1063, *p*=.93) in active lever presses nor any significant Condition × Sex two-way interactions (F_1,44_ =0.1385, *p*=.71), indicating that patterns of responding across training did not differ as a function of condition and sex. No significant day × sex (F_2.386,105_ = 1.621, *p*=.20) or day × condition (F_2.386,105_ = 2.43, *p*=.11) interactions were observed in active lever presses.

Main effects were examined next, and we found a significant main effect of Day in rewards (*F*_2.856,122.8_ = 9.19, *p*<.0001) and in active lever presses (*F*_2.386,105_ = 22.21, *p*<.0001), reflecting a change in rewards received and responding across training days. There were no main effects of condition (F_1,44_ =0.017, *p*=.89) or sex (F_1,44_ =0.6846, *p*=.41) in active lever pressing during FR1.

Motivation for social reward was assessed using a PR schedule. A two-way ANOVA revealed a significant main effect of condition across multiple measures of operant responding, including rewards earned (*F*_1,43_=6.512, *p*<.05), active lever presses (*F*_1,43_=8.069, *p*<.01) (Fig. 2C), and breakpoint (*F*_1,43_=8.013, *p*<.01) (Fig. 2D), with POE rats exhibiting reduced responding relative to controls across all measures. There were no significant main effects of sex in rewards (*F*_1,43_ <1.0, *p*=.63), active lever presses (*F*_1,43_ <1.0, *p*=.84), or breakpoint (*F*_1,43_<1.0, *p*<.70) and no significant condition × sex interactions, in rewards (*F*_1,43_<1.0, *p*=.63), active lever presses (*F*_1,43_ <1.0, *p*=.90), or breakpoint (*F*_1,43_ <1.0, *p*=.92), indicating that these effects were not sex dependent.

**Figure 2.**
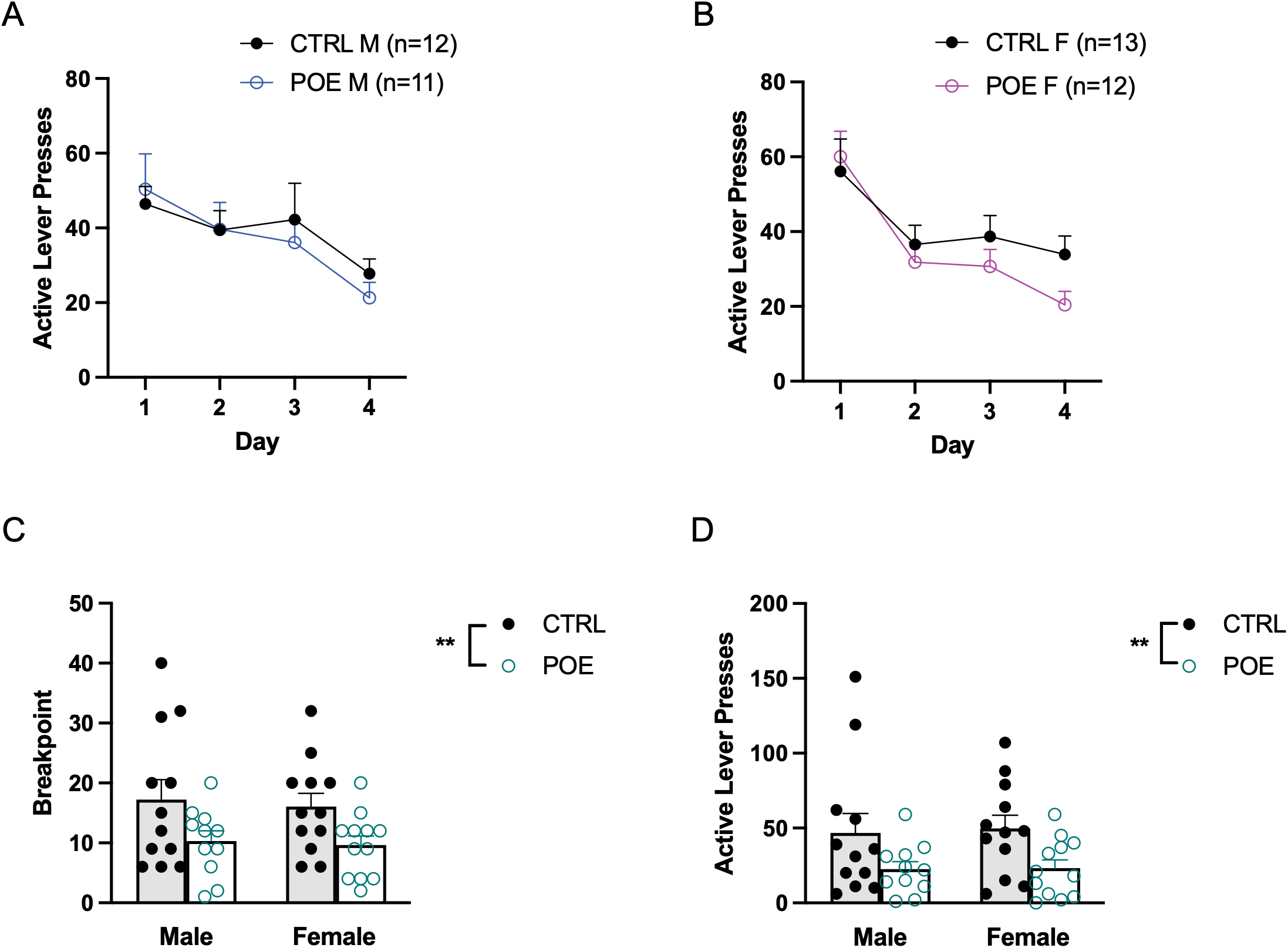
Effects of POE on social self-administration. A) Number of active lever presses on an FR1 schedule over 4 days in males. B) Number of active lever presses on an FR1 schedule over 4 days in females. C) Breakpoint ratio on a PR schedule. D) Number of active lever presses on a PR schedule. N=11-13/group. ** *p* <.01. Data are represented as means ± SEMs. CTRL, control; POE, Predator Odor Exposure; M, Male; F, Female.

## Discussion

Although POE has been shown to alter social behavior during development, particularly during the juvenile period, it remains unclear whether these effects persist into adulthood as enduring changes in the motivational value of social reward. Moreover, prior work has not clearly distinguished whether such behavioral alterations reflect impairments in task acquisition or performance versus selective changes in incentive motivation. An additional unresolved question is whether early disruptions in social behavior generalize to other forms of reward or remain specific to social behavior. The present study addresses these gaps by examining the impact of early life POE on adult responding for both sucrose reward and social interaction using operant self-administration paradigms. By assessing behavior under both FR and PR schedules, this approach allows for the dissociation of reward-directed responding and effort-based motivation while determining whether ELA produces domain-specific or generalized alterations in reward processing.

Importantly, POE did not alter the acquisition of social self-administration, as responding across FR schedules was similar across groups. Although responding varied across days, there were no effects of condition or sex during this phase, indicating that POE did not impact social self-administration on an FR1 schedule of reinforcement. These findings suggest that POE does not impair the ability to learn the action–outcome contingency or perform the operant response required to obtain social interaction. Clear group differences emerged under the PR schedule, where POE animals exhibited reduced breakpoint, active responding, and rewards earned relative to controls. Because PR is widely interpreted as an index of incentive motivation or willingness to work for a reinforcer^38,39^, these results indicate that POE reduces the motivational value of social interaction in adulthood. Altogether, this suggests that POE selectively impacts social motivation rather than the acquisition or performance of the operant response.

These findings extend prior work demonstrating that POE reduces juvenile social play^29^, a behavior highly sensitive to developmental experience^17,40,41^. The present results build on this work by showing that early disruptions in social behavior are not transient but instead persist into adulthood as diminished motivation to obtain social interaction. This continuity across developmental stages suggests that ELA may produce lasting changes in the valuation of social stimuli, potentially biasing behavior away from social engagement even when opportunities for interaction are available.

In contrast to social self-administration, no effects of POE were observed for sucrose motivation under the PR schedule, suggesting that effort-based motivation for sucrose reward remains largely intact following early-life threat exposure. Rather, effects of POE on sucrose responding were limited to the FR1 self-administration phase and were different across sex. Specifically, our POE females showed increased responding in comparison to control females or POE males. These findings do not support broad alterations in motivation for sucrose reward; however, the elevated responding observed during FR1 self-administration raises the possibility that POE may increase sensitivity to palatable reward or alter other aspects of hedonic processing in a sex-biased manner. In contrast to the reductions observed in social motivation, these findings suggest that the behavioral consequences of POE may differ across reward domains.

An important consideration when interpreting these findings is the specific dimension of adversity modeled by POE. Recent frameworks in the human adversity literature argue that experiences involving threat, deprivation, and unpredictability represent distinct dimensions of adversity that may influence development through partially dissociable mechanisms^42-44^. Much of the preclinical ELA literature has focused on paradigms such as maternal separation (MS) or limited bedding and nesting (LBN), which primarily manipulate caregiving environments and have been conceptualized as modeling aspects of deprivation or environmental unpredictability^10,45,46^. Because these paradigms alter the environment shared by the dam and offspring, developmental outcomes may reflect both direct effects on pups and indirect effects mediated through changes in maternal care. In contrast, POE exposes neonates directly to repeated threat-related cues during a sensitive developmental period, with exposure restricted to the offspring rather than including the dam. This feature distinguishes POE from many commonly used ELA paradigms and provides a unique opportunity to examine the effects of developmental threat exposure independent of alterations in the maternal environment.

Previous studies using MS and LBN have reported lasting changes in behavior. For example, LBN, a model characterized by resource scarcity and fragmented maternal care^46^, has been shown to reduce juvenile social play and induce broader alterations in reward processing, including anhedonia-like phenotypes^15,47^. Similarly, MS paradigms have been associated with long-lasting changes in social interaction and social approach behaviors^48-50^. We previously demonstrated that POE reduces juvenile social play behavior^29^, and the present findings extend these observations showing that these effects persist into adulthood as reduced motivation to obtain a social reward. Together, these findings suggest that direct developmental exposure to threat-related cues is sufficient to produce enduring alterations in social motivation. Williams and colleagues reported that early resource scarcity altered motivation for both social and sucrose rewards in a sex- and reinforcer-dependent manner^31^, highlighting that different forms of adversity may produce distinct effects on reward-related processes despite converging on similar social outcomes.

By examining responding for both social interaction and sucrose reward, the present study sought to determine whether the behavioral consequences of POE were specific to the social domain or reflected broader alterations in reward processing. The reduction in social motivation observed in our study suggests that developmental exposure to threat may have lasting consequences for social reward-related processes. Given that social behavior requires ongoing evaluation of environmental safety and social approach decisions^51,52^ , alterations in social motivation may represent a particularly enduring consequence of early-life threat exposure. This interpretation is further supported by our previous findings demonstrating reduced juvenile social play following POE, suggesting that developmental threat exposure disrupts social behavior across multiple stages of development. These findings suggest that developmental threat exposure may have enduring consequences for social reward-related processes. Future studies examining motivation and response patterns across additional reinforcers will be important for determining whether the effects of developmental threat exposure produce domain-specific deficits in social motivation or reflect broader alterations in reward valuation.

Reduced motivation for social interaction is a well-established feature of several stress-related psychiatric conditions, including depression and post-traumatic stress disorder (PTSD), characterized by social anhedonia, diminished reward processing, and reduced social engagement^53-55^. In clinical populations, disruptions in social reward processing, such as diminished responsiveness to positive social stimuli and decreased motivation to engage in social interactions, are linked to symptoms of anhedonia and social withdrawal^53,56^. Notably, ELA has been consistently linked with an increased risk for these disorders, as well as with long-term alterations in social behavior and affective functioning^4,57,58^. Behavioral manifestations of stress exposure often include reduced social engagement and avoidance of social interactions, behaviors that parallel the reduced motivation for social reward observed in our study. More broadly, social withdrawal and diminished social motivation are often linked to negative emotional outcomes. This underscores the importance of intact social reward processing for mental health. Our findings suggest that early exposure to threat-related cues may bias long-term behavioral trajectories by reducing the motivational value of social interaction, providing a potential behavioral mechanism through which ELA contributes to persistent social dysfunction and increased vulnerability to stress-related psychopathology.

## Acknowledgements

This work was supported by the National Institutes of Health (R21 DA059966, R01 DA049837, R01 DA056534, R34 DA061483, R21 DA062844, K99/R00 DA060266), the National Science Foundation (IOS-2313253, IOS-2208822), and the Wilczynski-Georgia Research Alliance Future Faculty Fellowship in Neuroscience. The content is solely the responsibility of the authors and does not necessarily represent the official views of the NIH or NSF.

